# To jump or not to jump: Comparing effects of phenotypic plasticity on the visual responses and escape behavior of locusts and grasshoppers

**DOI:** 10.1101/2025.08.26.672473

**Authors:** Soumi Mitra, Saina Namazifard, David M Bellini, Alexis Sarne, Bidisha Halder, Margaret R Eisenbrandt, Aliya Macknojia, Jiayi Luo, Sammie Xie, Hojun Song, Chenghang Zong, Fabrizio Gabbiani, Richard B Dewell

**Author notes:** co-senior authors (Chenghang Zong, Fabrizio Gabbiani, Richard B Dewell).

## Abstract

Locusts exhibit remarkable phenotypic plasticity changing their appearance and behavior from solitary grasshoppers to gregarious locusts when population density increases. These changes include morphological differences in the size and shape of brain regions, but little is known about plasticity within individual neurons and alterations in behavior not directly related to aggregation or swarming. We examined looming escape behavior and the properties of a well-studied collision-detection neuron in gregarious and solitarious animals of three closely related species, the desert locust (*Schistocerca gregaria*), the Central American locust (*S. piceifrons*) and the American bird grasshopper (*S. americana*). For this neuron, the lobula giant movement detector (LGMD), we examined dendritic morphology, membrane properties, gene expression, and looming responses. Gregarious animals reliably jumped in response to looming stimuli, but surprisingly solitarious desert locusts did not produce escape jumps. These solitarious animals also had smaller LGMD dendrites. This is the first study done on three different species of grasshoppers to observe the effects of phenotypic plasticity on the jump escape behavior, physiology and transcriptomics of these animals. Surprisingly, there were little differences in these properties between the two phases except for behavior. For all the three species, gregarious animals jumped more than solitarious animals, but no significant differences were found between the two phases of animals in the electrophysiological and transcriptomics studies. Our results suggest that phase change impacts mainly the motor system and that the physiological properties of motor neurons need to be characterized to understand fully the variation in jump escape behavior across phases.

New & Noteworthy (74 words): Swarming is observed in some grasshopper species, called locusts. We compared jump escape behavior between gregarious and solitarious grasshoppers and locusts, as well as LGMD responses to looming stimuli, and analyzed the morphological differences in this neuron. This study provides insights into the effects of phase change on the visual system of locusts and grasshoppers as it relates to looming-evoked jump escape behavior. In this context, our results suggest that phenotypic plasticity mainly impacts the motor system.

## INTRODUCTION

Phenotypic plasticity is defined as an organism’s ability to change its phenotype in response to environmental changes. The plasticity could be behavioral, morphological, developmental, physiological, or reproductive (Ghalambor et al., 2010; Simões et al., 2016; Song et al., 2017; Whitman & Agrawal, 2009). Desert locusts offer a remarkable example of phenotypic plasticity; in response to an increase in population density, cryptic solitarious animals turn into gregarious locusts of conspicuous color. The bi-directional, density-dependent switch between the solitarious and gregarious phases of locusts is known as phase polyphenism; this property has led to locusts being one of the most extensively studied animal models of phenotypic plasticity (Gotham & Song, 2013; Kilpatrick et al., 2019; Panhwar & Mustafa, 2022; Simões et al., 2016; Thompson, 1999). Plastic phenotypic changes are expected to be beneficial. For instance, a change in color helps solitarious locusts merge with their surroundings to avoid predation (Peralta-Rincon et al., 2017; Sword, 2002). In dry climatic regions, when food sources are plentiful after rainfall, the population of initially rare locusts increases until it reaches a point where vegetation becomes scarce due to foraging. This often leads to solitarious locusts (usually at densities of < 3 animals per 100 m^2^) encountering each other on resource patches thus triggering aggregation (reaching 100,000 animals per 100 m^2^). Solitarious individuals show significant gregarious characteristics within 4-8 hours of aggregation (Simões et al., 2016). The degree of phase polyphenism in grasshoppers of the *Schistocerca* genus differs between species, with some exhibiting pronounced changes, e.g., the desert locust (*S. gregaria*) and the Central American locust (*S. piceifrons*); no phase change, e.g., the gray bird grasshopper (*S. nitens*); or intermediate polyphenism, e.g., the American bird grasshopper (*S. americana*; Kilpatrick et al., 2019; Song et al., 2017; Sword, 2003)

Like other sighted animals, locusts and grasshoppers rely on vision to avoid collision or predation. Looming sensitive neurons that respond selectively to objects approaching on a collision trajectory have been described in several species, including pigeons (Sun & Frost, 1998; Xiao et al., 2006), vinegar flies (*Drosophila melanogaster*) (Ache et al., 2019; Allen et al., 2006; De Vries & Clandinin, 2012; Jang et al., 2022; Von Reyn & Card, 2012), zebrafish (Dunn et al., 2016; Fotowat & Engert, 2023; Marquez-Legorreta et al., 2020; Temizer et al., 2015), and grasshoppers (Gabbiani et al., 1999; McMillan & Gray, 2015; Rind & Simmons, 1999; Schlotterer, 1977). Postsynaptic to the Lobula Giant Movement Detector (LGMD), the descending contralateral movement detector (DCMD) neuron in grasshoppers is involved in triggering escape behavior (Burrows & Fraser Rowell, 1973; Fotowat et al., 2011). The LGMD is located in the optic lobe and synapses with the DCMD resulting in 1:1 spiking. The DCMD axon faithfully relays spike trains from the brain to the metathoracic ganglion where it synapses with the motoneurons that play a key role in jumping (Burrows & Fraser Rowell, 1973; O’Shea & Williams, 1974). Electrophysiological studies and modelling of the LGMD have produced detailed descriptions of its active membrane properties and their roles in neuronal computation (Dewell et al., 2022; Dewell & Gabbiani, 2018; Gabbiani et al., 2002; Peron et al., 2007; Peron & Gabbiani, 2009).

Previous research on the effect of phase change in desert locusts found gross anatomical density-dependent differences in the brain and optic lobe morphologies (Burrows et al., 2011; Ott & Rogers, 2010). Additionally, DCMD responses to looming stimuli were reported to be higher in gregarious desert locusts than isolated ones (Matheson et al., 2004; Rogers et al., 2007). Whether these phase differences result in differences for escape behavior has not been tested. We thus assessed the impact of density-dependent phase change on jump escape behavior and LGMD properties in species with varying degrees of polyphenism (*S. gregaria, S. piceifrons*, and *S. americana*; Fig 1).

**Figure 1.**
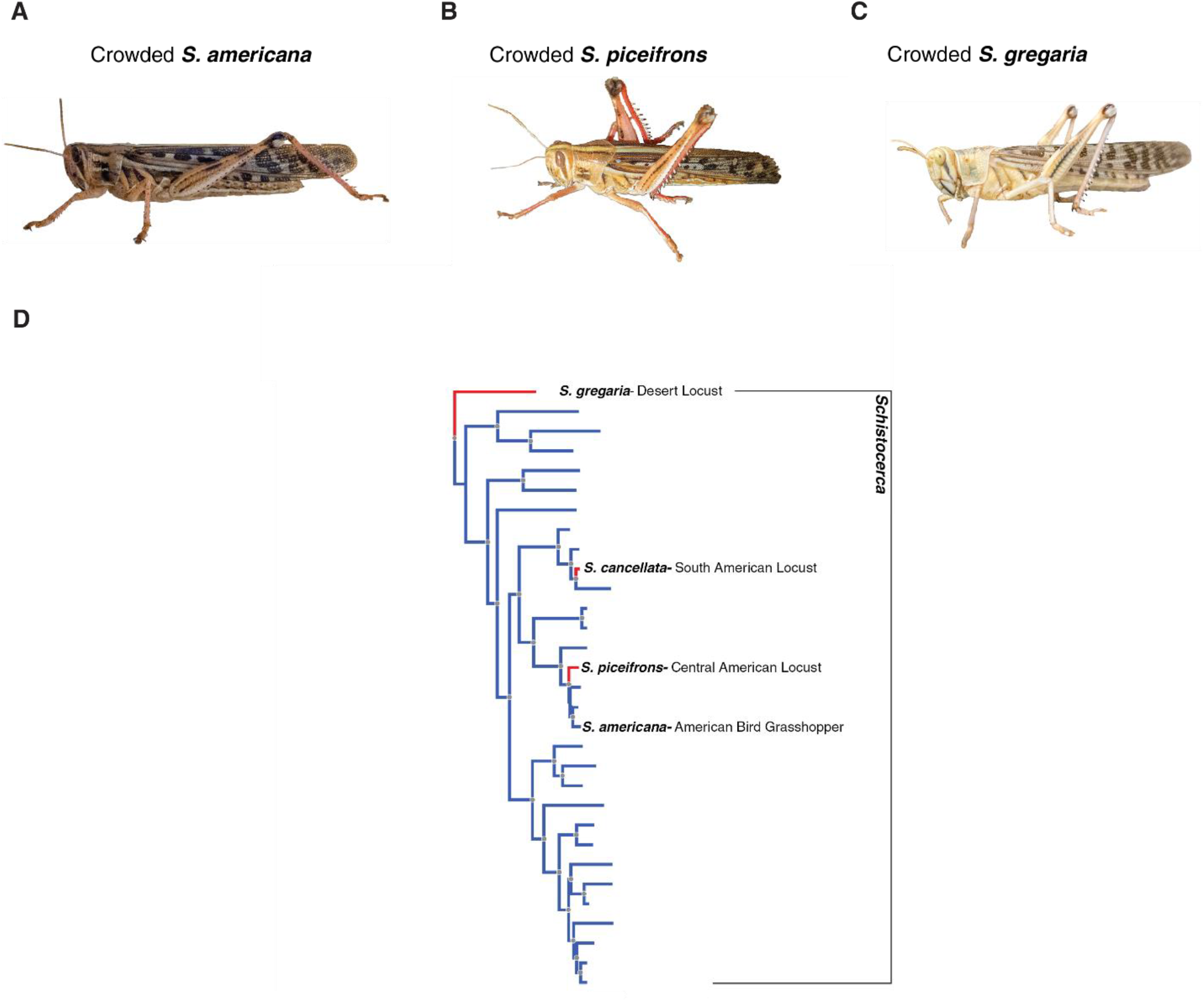
(A-C) Visible differences in coloration of the adults within the crowded (gregarious phase) animals belonging to *Schistocerca americana, S. gregaria* and *S. piceifrons*. (D) Bayesian phylogenetic tree showing that *S. piceifrons* (swarming species) and *S. americana* (non-swarming species) emerged from a lineage that shares a common ancestor with swarming *S. gregaria*. The red bars represent swarming species and illustrate that swarming has independently evolved multiple times in the *Schistocerca* genus. (Panel D adapted from Song et. al, 2017.)

Here we demonstrate a dramatic density-dependent plasticity in escape behavior, wherein gregarious animals of each species jumped more frequently than the solitarious ones. We then conducted a detailed study of the neuronal, physiological, anatomical, and transcriptional characteristics of the LGMD neuron within both phases of *S. gregaria* and *S. americana* in pursuit of the underlying causes of the behavioral difference. We couldn’t include *S. piceifrons* in these studies due to lack of availability of *S. piceifrons* at the time these experiments were conducted. Unlike previous investigation of *S. gregaria* (Matheson et al., 2004; Rogers et al., 2007), we did not find substantial differences in the DCMD firing between the two phases of either species. Furthermore, there were no differences in the LGMD dendritic morphology, membrane properties, or gene expression between phases that could account for the behavioral differences. These data suggest that phase-dependent difference in escape behavior is due to phenotypic plasticity of motor output rather than sensory processing.

## MATERIALS & METHODS

### Animals

All experiments were conducted using adults of both male and female grasshoppers (*S. americana*) and locusts (*S. gregaria, S. piceifrons*; 8-12 weeks old). Gregarious animals were reared in a dense colony, and solitarious animals were reared individually in small cages preventing them from touching, seeing, or smelling each other (Gotham & Song, 2013; Kilpatrick et al., 2019). Both groups were reared under 12:12-h light-dark conditions and had been in isolated or crowded conditions for at least two generations (Foquet, 2020). Animals were selected for health and size without randomization and investigators were not blinded to experimental conditions. Due to space constraints and legal restrictions on the use and maintenance of non-native locust species in the U.S., we had to limit the number of experimental animals. However, the number of replicates tested in this study is sufficient to draw adequate statistical conclusions.

### Visual stimuli

Visual stimuli were generated using MATLAB (RRID:SCR_001622; see https://scicrunch.org/resources for details) and the Psychophysics Toolbox (PTB-3, RRID:SCR_002881) on a personal computer. For DCMD experiments, stimuli were presented on a cathode ray tube monitor refreshed at 200 frame/s (LG Electronics, Seoul, Korea). For behavior experiments, a monitor operating at 240 frame/s was used (Acer Predator XB253Q). Looming stimuli are the two-dimensional projections of an object approaching on a collision course with the animal at constant velocity. Two different types of looming stimuli were used for jump experiments and extracellular DCMD recordings: black (OFF) and white (ON) presented on a background of opposite polarity. Looming stimuli consisted of a square simulating a solid object with half-size *l* approaching at constant speed *v* and were shown at the screen’s maximum contrast. As previously described, the expansion profile is characterized by the ratio *l/*|*v*| (in units of time; Gabbiani et al., 1999, Gabbiani et al., 2002)

### Jump experiments

Jump experiments were conducted using ON and OFF stimuli at three different *l/*|*v*| values: 20, 50 and 80 ms. These values were previously found to elicit reliable escape behavior. (Dewell & Gabbiani, 2018; Fotowat & Gabbiani, 2007; Gabbiani et al., 2001). Adult grasshoppers of both sexes were used for these experiments. Stimuli were shown to the animals in pseudo-random order, with at least a 10 min interval between two consecutive trials. 10 gregarious *S. piceifrons*, 9 solitarious *S. piceifrons*, 10 gregarious *S. gregaria*, 10 solitarious *S. gregaria*, 15 gregarious *S. americana* and 21 solitarious *S. americana* were used; each animal was shown up to 50 stimuli over up to 2 weeks of testing. Trials where the animal moved from the testing platform and did not align with the stimulus were excluded from analysis. Videos were recorded with a high-speed digital CMOS camera (BFS-U3-16S2M, Flir), equipped with a 12 mm lens (MVL12M23; Navitar). Image frames were recorded at 200 frames per second with the acquisition of each frame triggered by the visual stimulus with a USB to TTL cable (FTDI TTL-232R-5V-WE). Videos were made from 8-bit images and saved in lossless motion JPEG format using custom MATLAB code. The time when the animal jumped relative to collision was calculated based on the synchronization of the stimulus frames with camera frames.

### Electrophysiological experiments

The surgical procedures used in these experiments have been previously described (Dewell & Gabbiani, 2018; Gabbiani & Krapp, 2006; Jones & Gabbiani, 2012). For extracellular DCMD recordings, an adult animal was mounted ventral side up on a narrow plastic holder. The cuticle around the neck area was carefully removed with the help of forceps to expose the nerve cords. The animal was then secured on a stand in front of the monitor and a pair of Formvar-coated stainless steel wire hooks were placed on the ventral nerve cord between the suboesophageal and prothoracic ganglia to record extracellular DCMD responses to visual stimuli (Gabbiani et al., 2002). Both ON and OFF stimuli were used, and *l/*|*v*| values equal to 20, 40, 60 and 80 ms were shown to all the *S. gregaria*; *l/*|*v*| values equal to 20, 50 and 80 ms were shown to all *S. americana*. In total, 8 solitarious *S. gregaria*, 9 gregarious *S. gregaria*, 9 solitarious *S. americana* and 10 gregarious *S. americana* were used. A few drops of ice-cold locust saline were added intermittently during dissection and during visual experiments to retain moisture around the nerve cords. For this part of the study, we used only *S. gregaria* and *S. americana* and not *S. piceifrons* due to the unavailability of *S. piceifrons* at the time these experiments were conducted.

Intracellular LGMD recordings were done in discontinuous current-clamp mode using thin-walled borosilicate glass pipettes filled with a solution containing 1.0 M potassium acetate and 1.5 M KCl, yielding electrode resistances of 12-20 MΩ (pipette outer/inner diameter: 1.2/0.9 mm; WPI, Sarasota, FL; (Dewell & Gabbiani, 2018; Jones & Gabbiani, 2012). Step currents of +/-1, +/-2 and +/-5 nA were used. Membrane potential (*V*_*m*_) and current (*I*_*m*_) were low pass filtered with cut-off frequencies of 10 kHz and 5 kHz, respectively. Recordings were digitized at a sampling rate of at least 20 kHz. We used a single-electrode clamp amplifier capable of operating in discontinuous current-clamp mode (typically ∼3 kHz, Axoclamp 2B, Molecular Devices).

### LGMD Staining & Imaging

#### Animal Preparation

A mature locust was mounted on a plastic holder dorsal side up and immobilized by vacuum grease. Except for the right eye used for visual stimulation, the head and neck were bathed in ice-cold locust saline (Laurent & Davidowitz, 1994). The carapace on the head between the two eyes was partially removed to expose the head capsule where the two optic lobes and brain are located. The gut was completely removed. The muscles in the head capsule were also removed. The cuticle around the neck was removed leaving only two pairs of trachea and nerve cords. The right optic lobe was desheathed with the help of fine forceps (Dumont #5; FST, Switzerland). A metal holder was inserted underneath the right optic lobe to elevate and stabilize it during intracellular recordings (Gabbiani & Krapp, 2006).

#### Staining

The LGMD synapses with the DCMD which leads to a unique 1:1 spike pattern. This characteristic was used to identify the LGMD accurately while the spikes from DCMD were recorded extracellularly with hook electrodes placed on the ventral nerve cord. A sharp intracellular electrode (thin-walled borosilicate glass pipette, see above) was filled with ∼3 µL of solution of 1.0 M potassium acetate, 1.5 M KCl and 0.9 µL of 10 mM Alexa Fluor 594, hydrazide, (Thermo Fisher Scientific, Waltham, MA). The dye was injected into the LGMD for a few minutes by iontophoresis using a small negative current passing through the electrode (-1 nA).

#### LGMD dendritic morphology

For dendritic reconstructions, stained LGMDs were imaged using a custom built 2-photon excited fluorescence laser scanning microscope (Zhu & Gabbiani, 2018). Two-photon image stacks of the LGMD were acquired using ScanImage and processed using MATLAB for further analysis. The LGMD image files obtained post processing were then imported into neuTube software (Feng et al., 2015) for tracing followed by Sholl morphological analysis using Neuron Tree software (Cuntz et al., 2010). Next, compartmental modelling including passive properties was performed using NEURON software, as described in (Peron et al., 2007).

#### LGMD Passive Simulations

To estimate the unknown parameters R_m_, R_a_, and C_m_ by fitting current clamp data, the LGMD morphology was implemented in NEURON and used to simulate these data in a passive model. The time constant τ_m_ was obtained by fitting the initial exponential portion of the experimental membrane potential data. R_m_ and R_a_ were estimated by fitting the passive steady-state model output to the experimental data, using the least-squares method. Both R_a_ and R_m_ were assigned uniformly across all the segments in the NEURON model. C_m_ was calculated from the relation C_m_ = τ_m_ /R_m_.

### Single Cell RNA-Sequencing of LGMD soma

Cell bodies of the LGMD were extracted from the lobula using a suction pipette and stored in Trizol at -80 ^º^C after dissecting the grasshopper brain and staining with Alexa 594 fluorophore as described earlier. cDNA was obtained followed by single cell RNA-Seq of the genes expressed in the LGMD and the lobula using snapTotal-seq (Niu et al., 2024). Cell bodies were extracted from 8 solitarious *S. gregaria* and 7 gregarious *S. gregaria* for these experiments.

After sequencing, data was mapped to the *Schistocerca gregaria* genome using the STAR aligner (Dobin et al., 2013) and the count matrix generated using UMI-tools (Smith et al., 2017) to filter out reads with duplicate Unique Molecular Identifiers (UMIs). Then, the data was analyzed using edgeR, an R package for RNA-seq analysis (Robinson et al., 2010). The genes were tested for significant differential expression in edgeR using the quasi-likelihood F-test, and then the p-values were corrected for multiple testing using the Benjamini-Hochberg (BH) method.

### Data Analysis and Statistics

Data was analyzed using custom MATLAB code (The MathWorks, Natick, MA). Fisher’s exact test was performed to look for statistical significance in jump probability, while the Wilcoxon Rank Sum test was used to determine statistical significance of differences in the jump time values before collision. Jump probabilities were calculated based on the median unbiased estimator of a binomial response model. An ANOCOVA test was done to check for any statistically significant difference in the mean Time of Peak Firing (PKT) and instantaneous firing rate (IFR) values. A Kruskal-Wallis test was performed to check for significant differences in membrane input resistances. Estimates of the DCMD and motor neurons’ instantaneous firing rate were computed by convolving individual spike trains with a Gaussian function (width: 20 ms) as described earlier (Gabbiani et al., 1999).

## RESULTS

Here, we studied differences in escape behavior between long-term gregarious and solitarious phases of three different species of locusts and grasshoppers: *S. piceifrons, S. gregaria* and *S. americana. S. gregaria* and *S. piceifrons* are *bona fide* locust species, whereas *S. americana* exhibits an intermediate phenotype with some characteristics of swarming species, in spite of the fact that it is a grasshopper and does not swarm in its natural environment (Sword, 2003).

### Gregarious animals jumped more than solitarious ones

Solitarious animals had been reared separated which prevented them from touching, seeing, or smelling conspecifics for at least 2 generations; gregarious animals were reared for multiple generations in a high-density laboratory colony. For each behavioral trial, an unrestrained animal was shown a black looming stimulus once it walked on a platform and its eye was aligned with the center of the stimulus (Fig. 2A). Three different *l/*|*v*| values were used for this experiment where *l* is the half-length of the simulated approaching square, *v* is the approach velocity, and *θ* is the angular half-size of the looming stimulus subtended at the retina (Fig. 2B). Gregarious *S. piceifrons* jumped more for all stimuli than solitarious *S. piceifrons* (Fig. 2C; p_FE_ = 1.9•10^-6^, 3.2•10^-6^, 0.0003; Fisher’s exact test). Similarly, jump probabilities for gregarious *S. gregaria* at the three *l/*|*v*| values were higher than those of the solitarious *S. gregaria* (Fig. 2D). Only one of the ten tested solitarious *S. gregaria* jumped during a loom – in response to one stimulus with *l/*|*v*| of 80 ms; none of the other nine solitarious *S. gregaria* animals jumped during a loom resulting in different jump probabilities for solitarious vs. gregarious animals (p_FE_ = 8.16•10^-8^, 2.92•10^-8^, 2.27•10^-7^). For *S. americana*, the jump probabilities were also higher for gregarious animals at all the three *l/*|*v*| values (p_FE_ = 1.1•10^-7^, 0.002, 0.0071). We also compared the timing of escape jumps relative to the time of projected collision. Gregarious *S. piceifrons* jumped earlier than the solitarious ones for each stimulus (Fig. 2F; p_WRS_ = 0.034, 0.044, 0.007; Wilcoxon Rank sum test). Gregarious *S. americana* jumped later than solitarious ones for the two faster looming stimuli (Fig. 2H; p_WRS_ = 0.0039, 0.046, 0.146). Long-term gregarious animals jump more than solitarious animals for every stimulus tested, but no consistent differences in jump timing were found across species.

**Figure 2.**
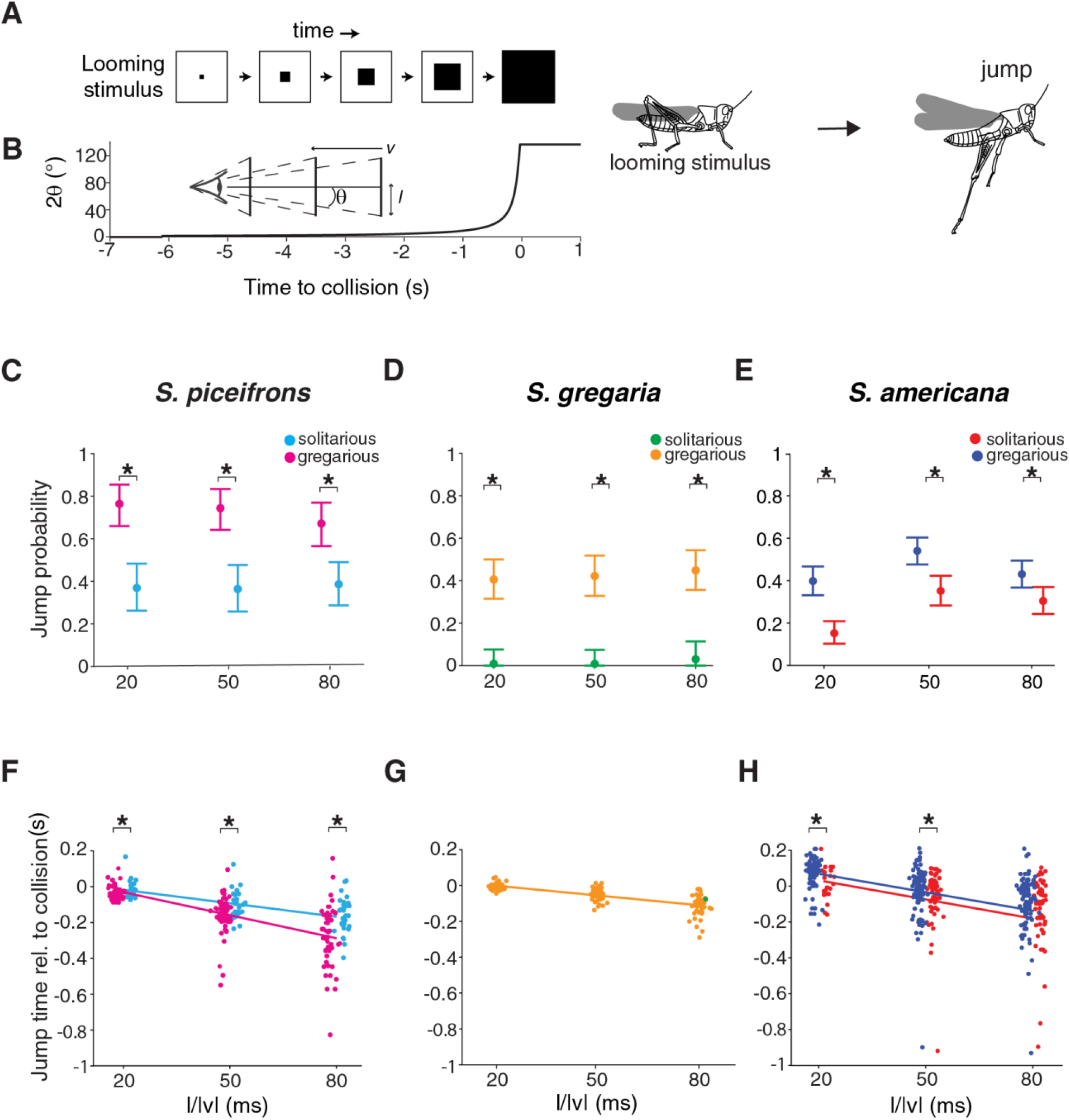
Jump probabilities were lower for solitarious locusts and grasshoppers. (A) Schematic illustration of looming stimulus. (B) Time course of loom’s angular size illustrating its non-linear progression during a jump experiment (*l/*|*v*|=80 ms; *l*=half length, *v*=velocity). (C-E) Jump probability of locusts (*S. piceifrons*, solitarious n = 9, gregarious n = 10; *S gregaria*, solitarious n = 10, gregarious = 10) & grasshoppers (*S. americana*, solitarious n = 21, gregarious n = 15). Gregarious locusts jump more to looming stimuli. Error bars are 95% confidence intervals. (F-H) Linear fit of jump times before collision. * Indicates p < 0.05, Fisher’s exact test (C-E) and Wilcoxon rank-sum test (F-H).

### Neural activity did not differ between phases

Looming-evoked jumps are triggered by descending activity of the DCMD neuron (Dewell et al., 2022; Dewell & Gabbiani, 2018; Fotowat et al., 2011), so we examined DCMD looming responses in the two phases of *S. gregaria* and *S. americana*. DCMD activity was recorded by placing a pair of wire hooks on the nerve cord (Fig. 3A). White and black looming stimuli of multiple half-size to speed ratios were presented to the contralateral eye, DCMD spikes were identified by spike thresholding, and instantaneous firing rates (IFR) were calculated (Fig. 3B). The peak IFR in response to black and white looms were measured (Fig. 3C-F). Only the *S. americana* responses to black looms with *l*/|*v*| of 80 ms were different between phases, with gregarious animals showing higher firing than solitarious ones (p_WRS_ = 0.002; Fig.3E). In these experiments we were diligent in dishabituating the animals between stimuli. This differs from previous findings in which DCMD IFR were higher in gregarious *S*.*gregaria* following greater habituation of solitarious animals (Matheson et al., 2004; Rogers et al., 2010).

**Figure 3.**
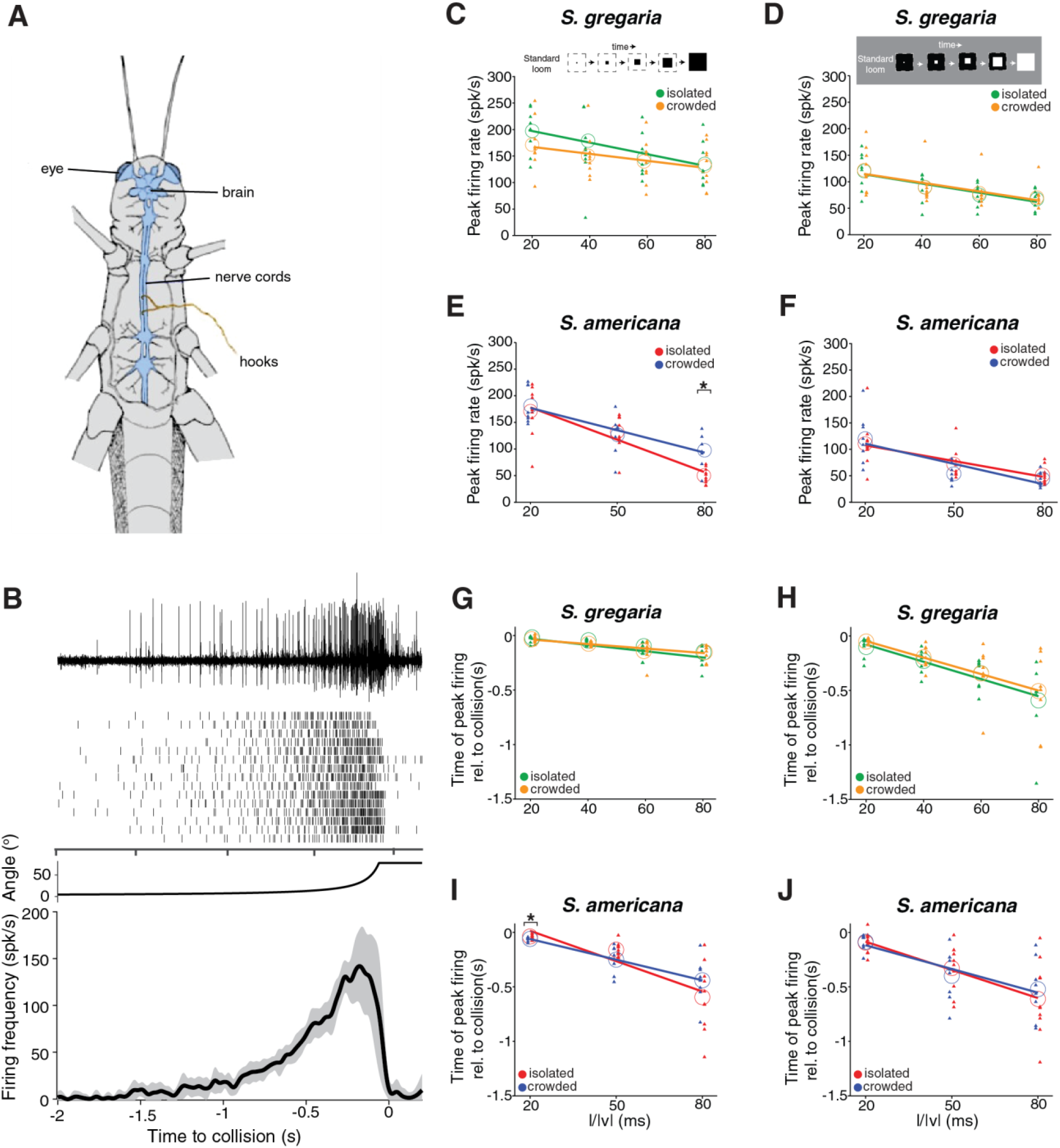
Comparison of DCMD responses between phases of locusts and grasshoppers. (A) Illustration of DCMD recording with wire hooks on the grasshopper’s ventral nerve cord. (B) Example recording and spike rasters of DCMD (top) from a solitarious *S. americana* animal, stimulus angular size (middle) and mean ± s.d. of DCMD instantaneous firing rate (IFR; bottom) for an example animal response to looms with *l*/|*v*| = 50 ms (C, D) Peak IFR to black & white looms in solitarious and gregarious *S. gregaria* at four different *l/*|*v*| values. Triangles show individual means; empty circles show population means and lines represent linear fit of individual means. (E, F). Peak IFR to black & white looms in isolated and crowded *S. americana* at three different *l/*|*v*| values, plotted as in C-D. (G, H) Time of the peak firing relative to collision for black (G) and white (H) looms in solitarious and gregarious *S. gregaria* at four *l/*|*v*| values. (I, J) Peak IFR timing of *S. americana*, plotted as in G and H. Solitarious *S. americana* had earlier responses for black looms with *l*/|*v*| = 20 and 80 ms; * indicate p <0.05, Wilcoxon rank sum test. Solitarious *S. gregaria* n = 8, gregarious n = 9, solitarious *S. americana* n = 10, gregarious n = 8.

We also compared the response timing for each stimulus (Fig. 3G-J). No differences were found in the time of peak firing between gregarious and solitarious *S. gregaria* (Fig. 3 G, H). For *S. americana*, the peak time of black loom responses was later for gregarious animals, with looms at an *l/*|*v*| value of 20 ms occurring after collision (p_WRS_ = 0.043; Fig. 3I). This is consistent with the behavioral responses to black looms which also showed later jumps in the gregarious *S. americana* (Fig. 2H). The response timing to white looms did not differ for solitarious and gregarious *S. americana*. Overall, our data showed that the DCMD responses between the phases of *S. gregaria* and *S. americana* were like each other.

### LGMD morphology is the same in solitarious and gregarious animals

So far, we observed that gregarious animals jumped more than the solitarious ones, but DCMD activity was similar between the two phases. Gross anatomy of the brains and optic lobes are different between phases of *S. gregaria* (Ott & Rogers, 2010), but individual neuron morphology has not been compared. We examined whether phenotypic plasticity has any significant effect on the morphology of the LGMD. To answer this question, we stained the LGMD by intracellular injection of fluorescent dye in both phases of *S. gregaria* and gregarious *S. americana* (Fig. 4). The LGMD has three distinct dendritic fields with the largest, field A, receiving retinotopically mapped excitation (Dewell et al., 2022; Peron & Gabbiani, 2009; Zhu & Gabbiani, 2016). The median size of field A when measured medial-laterally in gregarious and solitarious *S. gregaria* was 230 µm (p_WRS_ = 0.56), the median size of field A in gregarious *S. americana* was ∼200 µm (p_WRS_ = 0.002; Fig. 4B). Similarly, for the dorso-ventral measurements, the median size of field A in *S. gregaria* was ∼460 µm for both phases (p_WRS_ = 0.40), while that of gregarious *S. americana* was smaller (p_WRS_ = 0.040). The 3-axis maximum projections of 2-photon stacks for selected example LGMDs show that the LGMD in *S. americana* is slightly more compact, with similar morphology as those of *S. gregaria* across phases (Fig. 4D). To further characterize the LGMD morphology, we performed Sholl analysis on the dendritic branching within field A (Bird & Cuntz, 2019). This analysis examines the number of dendritic branches within each radial distance from the base of the dendritic field (Fig. 4E). Like with the 2-D measures (Fig. 4B, C), the Scholl analysis showed no difference in dendritic morphology between phases of *S. gregaria* (p_KS_ = 0.85, Kolmogorov-Smirnov test), but that the gregarious *S. americana* LGMD was more compact than either (p_KS_ = 0.04, 0.02).

**Figure 4.**
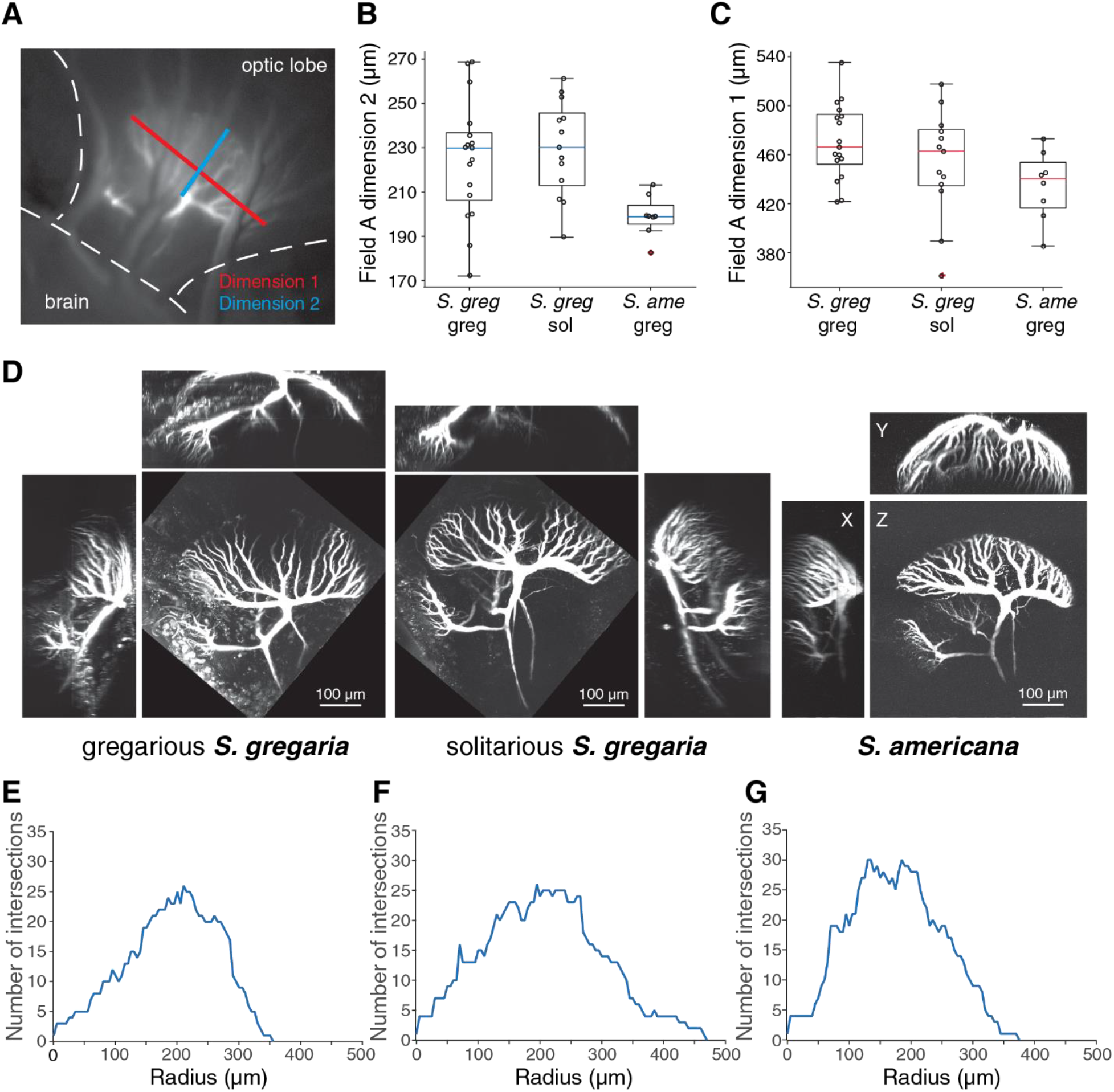
LGMD morphology of the two phases. (A) CCD image of an LGMD stained with Alexa 594. The size of the dendritic field was measured in two different dimensions: medial-lateral (dimension 1, red) and dorso-ventral (dimension 2, blue). (B, C) For both dimensions, *S. americana* LGMD was smaller (p = 0.002 and p = 0.040; Wilcoxon rank sum), no significant differences in LGMD size between gregarious and solitarious *S. gregaria* (p = 0.56 and p = 0.40), empty black circles represent individual data points. (13 solitarious *S. gregaria*, 17 gregarious *S. gregaria*, 8 gregarious *S. americana*). (D) 2 photon stacks of the LGMD showing smaller LGMD in *S. americana* (right). (E) Sholl analysis of LGMD of gregarious *S. americana* (right), solitarious *S. gregaria* (center) and gregarious *S. gregaria* (left) showed that the distribution of intersections was significantly different in gregarious *S. americana* (Kolmogorov-Smirnov test, p value <0.05).

### LGMD input resistance was consistent between the two phases

To test whether the membrane properties in the LGMD differ between the phases, we conducted *in vivo* dendritic intracellular recordings. For these experiments, we injected both hyperpolarizing and depolarizing step currents into the LGMD and measured the change in membrane potential to calculate input resistance (Fig. 5A-B). The membrane input resistance (R_i_) was measured in the LGMD for 4 solitarious and 5 gregarious *S. americana* and 3 gregarious *S. gregaria*. The median R_i_ was compared for each of the 6 input currents between the 3 groups with no difference at any current (p > 0.6, Kruskal-Wallis test; Fig. 5B). Like with the morphology of the neuron (Fig. 4), the electrotonic size of the neuron did not differ between phases.

**Figure 5.**
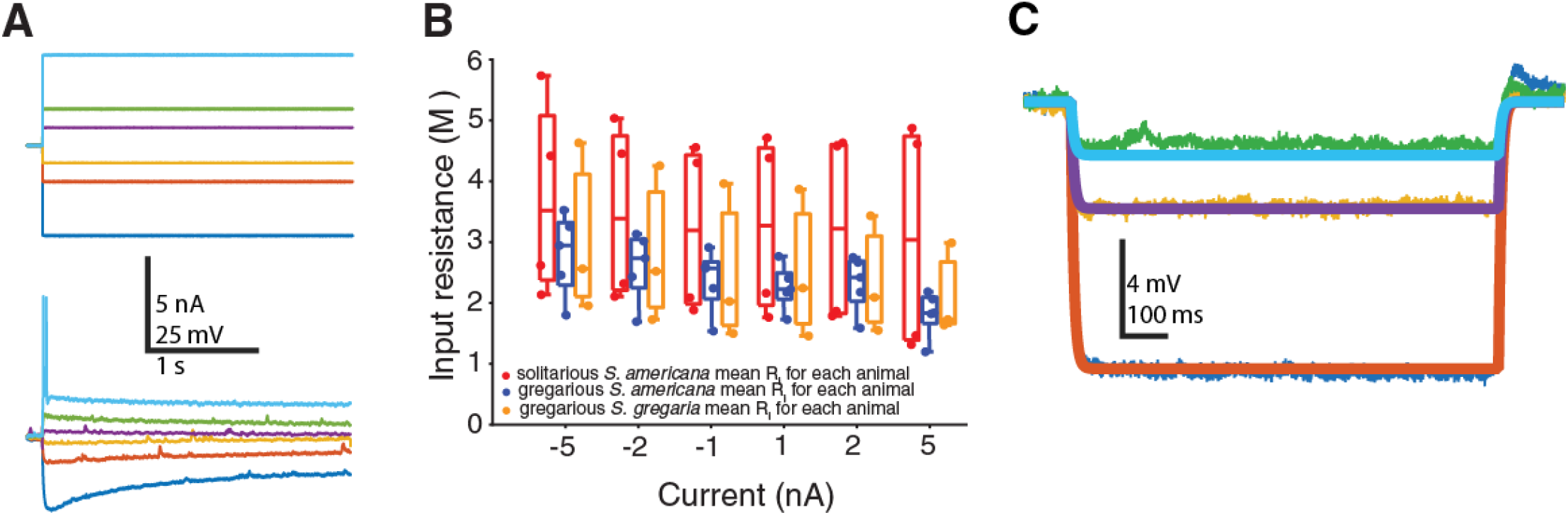
Measuring differences in LGMD membrane input resistance (R_in_) to step currents. (A) Recorded I_M_ and V_M_ traces showing step currents (-5, -2, -1, 1, 2 and 5 n A) injected to the LGMD and voltage change generated due to current injection. (B) Comparison of LGMD membrane input resistance in solitarious (n = 4), gregarious (n = 5) *S. americana* and gregarious *S. gregaria* (n = 3). Box plot shows the median, quartiles, and data extents of R_in_ value for each group; dots show mean (R_in_) value for each animal. None of the (R_in_) values in the three groups were statistically significant (Kruskal Wallis test). (C) Example Vm traces and best fits of the passive model (injected currents: -1, -2 and -5 nA).

### LGMD Passive Compartmental Simulations

Comparing membrane properties of the LGMD upon injecting current *in silico* showed no significant differences in the fitted specific axial resistance (R_a_), membrane resistance (R_m_) and membrane capacitance (C_m_) of the neuron in three animals from three different groups of grasshoppers. While the axial resistance of the LGMD in gregarious *S. americana* was 72.3 Ω cm, the same in gregarious *S. gregaria* was 61.2 Ω cm and 75.0 Ω cm in solitarious *S. gregaria*. The membrane resistances were quite close to each other, ranging from 5.9 kΩ cm^2^ in gregarious *S. americana* to 4.6 kΩ cm^2^ and 6.3 kΩ cm^2^ in gregarious and solitarious *S. gregaria* respectively. The C_m_ in gregarious *S. americana* was 1.1 µF/cm^2^, while for gregarious and solitarious *S. gregaria* it amounted to 1.4 µF/cm^2^ and 1.1 µF/cm^2^, respectively. These values match with the ones shown in a previous study done on *S. americana* (Peron et al., 2007). In general, the simulated neuron models accurately fitted the experimental data (Fig. 5C). The average R^2^ value for the traces was 0.9.

### Gene expression in the LGMD was consistent between phases

Individual LGMD somata were extracted from *S. gregaria* of both phases and the total mRNA of the cells were sequenced (Niu et al., 2024). In each cell, RNA reads of 13,000-18,000 unique genes were detected. Of these genes, only one gene was differentially expressed between phases, with greglin (LOC126331978) more highly expressed in the LGMD of gregarious *S. gregaria* compared to solitarious animals (Fig. 6A). For each LGMD that was harvested for sequencing, the rest of the lobula was used for a bulk RNA sequencing control. There were only six differentially expressed genes between the phases in the bulk controls, with two vitellogenin genes highly expressed in the lobula of gregarious *S. gregaria* (Fig. 6B). This shows surprisingly little change in gene expression of visual neurons between phases.

**Figure 6.**
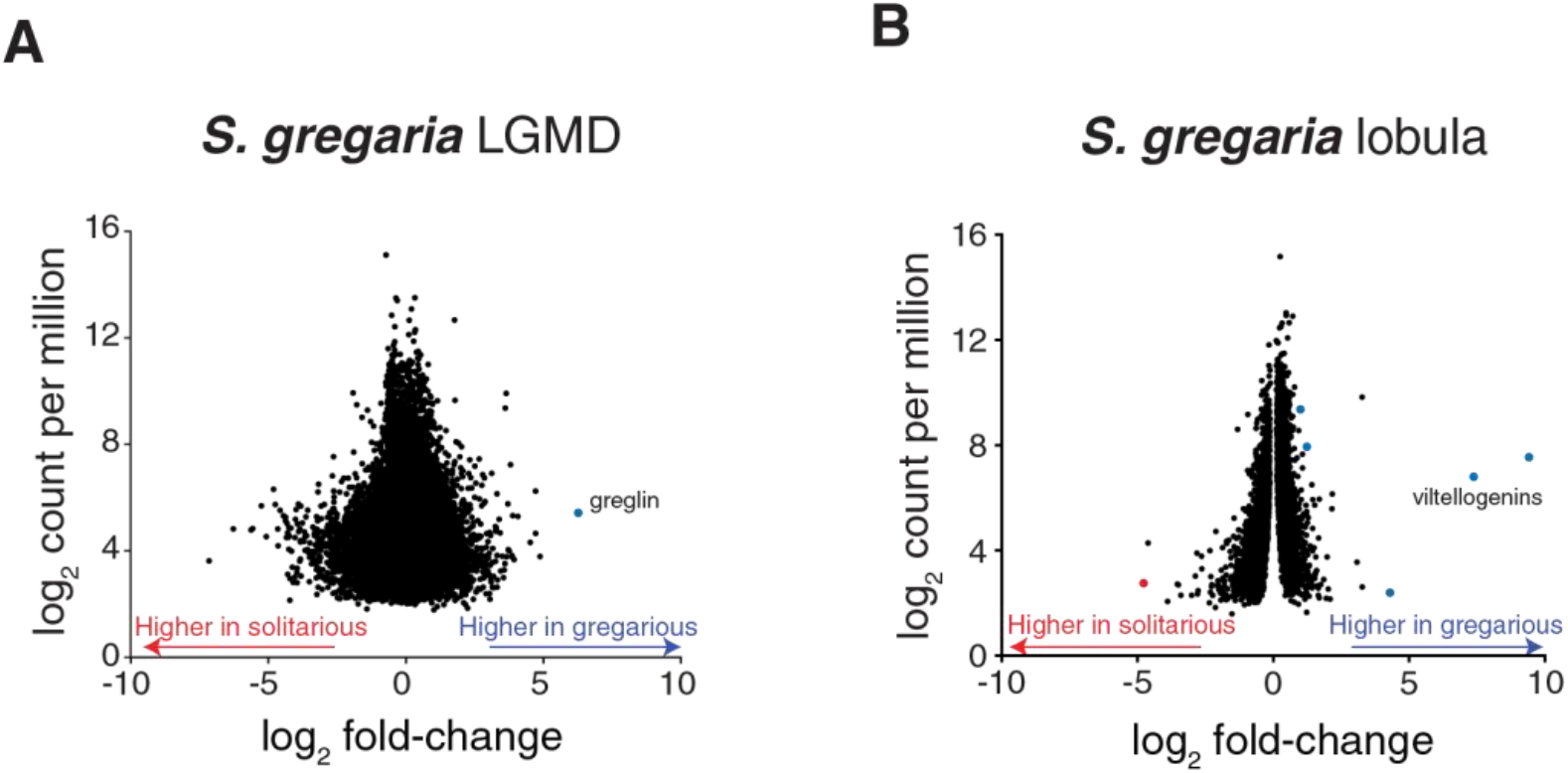
Differential gene expression between two phases of *S. gregaria*. (A) Analysis of single cell RNA-seq data revealed only one significantly differently expressed gene (DEG), greglin, within the LGMD. (B) Similarly, among all the genes expressed in the lobula that were analyzed, only six DEGs were found: vitellogenins (LOC126298903 & LOC126298904), trichohyalin (LOC126335968), solute carrier family 2 facilitated glucose transporter (LOC126354871), and an uncharacterized protein (LOC126278381) were higher in gregarious animals, and estradiol 17-beta-dehydrogenase 2-like (LOC126341510) was higher in solitarious animals.

## DISCUSSION

We conducted a study wherein we tested for density-dependent differences in looming escape jump responses of *S. piceifrons, S. gregaria* and *S. americana* reared under either solitarious or gregarious conditions. We found that for all three species, the gregarious animals jumped significantly more than their solitarious counterparts (Fig. 2). This study is the first to examine phase differences in looming-evoked escape and the first focusing on a possible phase-dependence in jumping or loom responses in *S. americana* or *S. piceifrons*. As previous studies found the responses of the LGMD to determine looming-evoked behavior, we followed up on these behavioral observations with an examination of the physiology, genetics, and morphology of the LGMD between phases. Surprisingly, we found no differences that could explain the phase-dependence of loom escape jumps.

Phenotypic plasticity results in behavioral, morphological and genetic changes in animals that are mediated by changes in environmental conditions (Agrawal, 2001; Burrows et al., 2011; Price et al., 2003; Uvarov, 1966). Earlier studies showed that the DCMD firing between the two phases of *S. gregaria* locusts varied significantly (Matheson et al., 2004; Rogers et al., 2007). The visual system, the motor system, and the musculature have been widely studied in grasshoppers and locusts (Gabbiani & Krapp, 2006; Matheson et al., 2004; Rogers et al., 2016). Until now, not much information was available about the differences in escape behavior, electrophysiological and morphological characteristics of the LGMD upon phase change in *S. gregaria* and *S. americana*. This sets the stage for our study and for a side-by-side comparison of the differences in escape behavior, DCMD firing patterns, LGMD shape and size, as well as differential gene expression within the LGMD between phases and species of grasshoppers and locusts. Currently, the only available data for direct comparison to our findings comes from *S. gregaria*.

Indeed, previous studies of *S. gregaria* found that solitarious grasshoppers have more muscular hind legs that help them jump faster and further than gregarious locusts (Rogers et al., 2016). According to Rogers et al., 2016, solitarious animals spent twice the energy per jump relative to gregarious ones. This energetic difference could be the reason why our solitarious animals jumped significantly less than gregarious ones. Due to the dearth of alternative explanations for these differences in escape behavior, we suggest that energy conservation in solitarious grasshoppers is a plausible explanation. Besides, we know that in the solitarious stage grasshoppers are cryptically colored helping them blend easily with their surroundings thus increasing their chances of survival by avoiding predation. In contrast, gregarious animals can be brightly colored, and easily visible. This could be another possible reason why isolated animals are less prone to escape jumping.

Looming escape jumps depend on the firing pattern of the LGMD neuron (Dewell et al., 2023; Fotowat et al., 2011), so we hypothesized that the difference in jump behavior between phases could be an outcome of differences in LGMD activity. To test this hypothesis, we took advantage of the 1:1 firing pattern of the LGMD and its post-synaptic neuron, the DCMD (O’Shea & Williams, 1974). Here we found that the DCMD firing patterns in the two phases of both species were not significantly different from each other and that they are unlikely to explain the differences in jump behavior (Fig. 3). Matheson et al, 2004 conducted the first study comparing DCMD firing rates in the two phases of *S. gregaria* and showed a clear difference between gregarious and solitarious animals (see also Rogers et al., 2007). This contrasts with our findings suggesting no such differences. These studies presented stimuli with short intertrial intervals magnifying the effect of habituation on DCMD firing rates across phases. In contrast, we paid particular attention to dishabituating the animal after each loom, by using, e.g., high frequency sounds, flashing lights, and tactile stimulation. Further, Gaten et al., 2012 noticed no differences in response between phases when waiting an hour between looms which would prevent any habituation, in agreement with our findings.

Since earlier studies have shown that gregarious *S. gregaria* brains are significantly larger than long-term solitarious ones (Burrows et al., 2011; Ott & Rogers, 2010) we compared the morphology of the LGMD between the two phases. There has been no previous comparison of individual cell morphology between phases, though. Our analysis of LGMD dendritic morphology in gregarious *S. americana*, solitarious and gregarious *S. gregaria* showed no significant differences in the dendritic arbors between the two phases of *S. gregaria*. Yet, the dendritic arbor of *S. americana* was smaller than that of *S. gregaria* (Fig. 4).

Having established that there was no difference in the shape or size of the LGMD, we wanted to determine if there were any differences in the membrane properties of the LGMD between the two phases. To answer this question, we recorded intracellularly from field A dendrites *in vivo* and measured membrane input resistance and resting membrane potential. Once again, our data showed that there were no significant differences in the properties of the LGMD between the two phases (Fig. 5). Since neither morphology nor input resistance were different, this suggests that the electrotonic characteristics of the LGMD are also consistent across phases. It is not possible to directly measure the axial resistivity, membrane resistance, or membrane capacity from these recordings, so we used previously described methods for estimating these properties from current steps using biophysical modeling (Peron et al., 2007). These simulations involved creating multi-compartmental models from the detailed dendritic morphologies of both phases (Fig. 4) and fitting the current clamp data (Fig. 5). The resulting simulations suggested consistent membrane properties across species and phase, with values similar to those found previously for *S. americana* (Peron et al., 2007).

Previous experiments on the active membrane properties of the LGMD have found specific ion channels to play critical roles in processing of looming stimuli (Dewell & Gabbiani, 2018; Peron & Gabbiani, 2009). We wanted to see if any receptor or voltage-gated ion channels present in the cell membrane are differentially expressed upon phase change. So, we extracted LGMD somata for single-cell RNA sequencing and found that only greglin was upregulated in gregarious *S. gregaria*. Greglin is a serine protease inhibitor induced by juvenile hormone that is involved in vitellogenesis (egg yolk production in vertebrates and invertebrates) (Derache et al., 2012; Guo et al., 2019). Absence of greglin results in immature oocytes and reduced egg numbers (Ciudad et al., 2006). In *S. piceifrons*, RNA sequencing of head tissue found greglin to be upregulated in gregarious locusts (Foquet, 2020). The connection between greglin and vitellogenesis is notable as there was increased expression of vitellogenins in the surrounding lobula (Fig. 6B). In addition to its role in egg production, vitellogenin is also involved in immune function and energy metabolism with different expression between castes in the brain of multiple social insect species (Hawkings & Tamborindeguy, 2025; Kohlmeier et al., 2018; Manfredini et al., 2018; Münch et al., 2015). The LGMD is an energetically expensive neuron and the reduced expression of greglin and vitellogenin in solitarious animals might be due to an energy saving strategy.

Gregarious and solitarious locusts differ in coloration, size, and behavior (Song et al., 2017). Additionally, we found clear differences in escape behavior for both locusts (*S. gregaria* and *S. piceifrons*) and a non-swarming grasshopper (*S. americana*). Despite this we found no differences in morphological, electrophysiological, or genetic properties of the LGMD between the phases that could explain the behavioral difference. This raises the question of where the phase-dependent difference arises and what other neurons might underly the change in behavior. The most likely candidates would be the jump coordination circuitry within the meta-thoracic ganglion. The DCMD synapses directly onto flexor and extensor motor neurons generating jumps and activity in these neuron differ between phases of *S. gregaria* (Burrows & Fraser Rowell, 1973; Rogers et al., 2007, 2016). Phase-dependent neuromodulatory changes of this circuitry depressing the synaptic connection from the DCMD onto these neurons could lead to reduced motor activity.

## Acknowledgements

We are grateful to collaborators at Texas A&M University for providing the isolated and crowded *S. gregaria* and *S. piceifrons* used here. Their locust husbandry made the present study as comprehensive as feasible. We thank Dr. Gregory Sword and Dr. Maeva Techer for providing valuable feedback on our manuscript. We thank Richelle Marquez for providing us with animals. We also thank Eleni Nasiotis for the cartoon drawing of grasshopper nerve cords in Fig. 3. This study was funded by NSF DBI 2021795 and NSF IOS-2212750.

## Data availability

All the data is available on Mendeley database

https://data.mendeley.com/preview/gxrwdk2nvt?a=10c161ef-fd7a-4351-a256-f45cfe145a70

## Notes

### Competing Interest Statement

The authors have declared no competing interest.

## REFERENCES

Ache, J. M., Polsky, J., Alghailani, S., Parekh, R., Breads, P., Peek, M. Y., Bock, D. D., Von Reyn, C. R., & Card, G. M. (2019). Neural basis for looming size and velocity encoding in the Drosophila giant fiber escape pathway. Current Biology, 29(6), 1073–1081.

Agrawal, A. A. (2001). Phenotypic plasticity in the interactions and evolution of species. Science, 294(5541), 321–326.

Allen, M. J., Godenschwege, T. A., Tanouye, M. A., & Phelan, P. (2006). Making an escape: Development and function of the Drosophila giant fibre system. 17(1), 31–41.

Bird, A. D., & Cuntz, H. (2019). Dissecting sholl analysis into its functional components. Cell Reports, 27(10), 3081–3096.

Burrows, M., & Fraser Rowell, C. H. (1973). Connections between descending visual interneurons and metathoracic motoneurons in the locust. Journal of Comparative Physiology, 85(3), 221–234. 10.1007/BF00694231

Burrows, M., Rogers, S. M., & Ott, S. R. (2011). Epigenetic remodelling of brain, body and behaviour during phase change in locusts. Neural Systems & Circuits, 1(1), 11. 10.1186/2042-1001-1-11

Ciudad, L., Piulachs, M., & Bellés, X. (2006). Systemic RNAi of the cockroach vitellogenin receptor results in a phenotype similar to that of the Drosophila yolkless mutant. The FEBS Journal, 273(2), 325–335.

Cuntz, H., Forstner, F., Borst, A., & Häusser, M. (2010). One Rule to Grow Them All: A General Theory of Neuronal Branching and Its Practical Application. PLoS Computational Biology, 6(8), e1000877. 10.1371/journal.pcbi.1000877

De Vries, S. E., & Clandinin, T. R. (2012). Loom-sensitive neurons link computation to action in the Drosophila visual system. Current Biology, 22(5), 353–362.

Derache, C., Epinette, C., Roussel, A., Gabant, G., Cadene, M., Korkmaz, B., Gauthier, F., & Kellenberger, C. (2012). Crystal structure of greglin, a novel non-classical K azal inhibitor, in complex with subtilisin. The FEBS Journal, 279(24), 4466–4478.

Dewell, R. B., Carroll-Mikhail, T., Eisenbrandt, M. R., Mendoza, A. F., Halder, B., Preuss, T., & Gabbiani, F. (2023). Convergent escape behaviour from distinct visual processing of impending collision in fish and grasshoppers. The Journal of Physiology, 601(19), 4355–4373. 10.1113/JP284022

Dewell, R. B., & Gabbiani, F. (2018). Biophysics of object segmentation in a collision-detecting neuron. eLife, 7, e34238. 10.7554/eLife.34238

Dewell, R. B., Zhu, Y., Eisenbrandt, M., Morse, R., & Gabbiani, F. (2022). Contrast polarity-specific mapping improves efficiency of neuronal computation for collision detection. eLife, 11, e79772. 10.7554/eLife.79772

Dobin, A., Davis, C. A., Schlesinger, F., Drenkow, J., Zaleski, C., Jha, S., Batut, P., Chaisson, M., & Gingeras, T. R. (2013). STAR: ultrafast universal RNA-seq aligner. Bioinformatics, 29(1), 15–21.

Dunn, T. W., Gebhardt, C., Naumann, E. A., Riegler, C., Ahrens, M. B., Engert, F., & Del Bene, F. (2016). Neural circuits underlying visually evoked escapes in larval zebrafish. Neuron, 89(3), 613–628.

Feng, L., Zhao, T., & Kim, J. (2015). neuTube 1.0: A new design for efficient neuron reconstruction software based on the SWC format. Eneuro, 2(1).

Foquet, B. (2020). Locust Phase Polyphenism in a Phylogenetic Framework: From Gene Expression to Behavioral Changes [Texas A&M University]. https://hdl.handle.net/1969.1/193004

Fotowat, H., & Engert, F. (2023). Neural circuits underlying habituation of visually evoked escape behaviors in larval zebrafish. Elife, 12, e82916.

Fotowat, H., & Gabbiani, F. (2007). Relationship between the Phases of Sensory and Motor Activity during a Looming-Evoked Multistage Escape Behavior. The Journal of Neuroscience, 27(37), 10047–10059. 10.1523/JNEUROSCI.1515-07.2007

Fotowat, H., Harrison, R. R., & Gabbiani, F. (2011). Multiplexing of Motor Information in the Discharge of a Collision Detecting Neuron during Escape Behaviors. Neuron, 69(1), 147–158. 10.1016/j.neuron.2010.12.007

Gabbiani, F., & Krapp, H. G. (2006). Spike-frequency adaptation and intrinsic properties of an identified, looming-sensitive neuron. Journal of Neurophysiology, 96(6), 2951–2962.

Gabbiani, F., Krapp, H. G., Koch, C., & Laurent, G. (2002). Multiplicative computation in a visual neuron sensitive to looming. Nature, 420(6913), 320–324. 10.1038/nature01190

Gabbiani, F., Krapp, & Laurent. (1999). Computation of Object Approach by a Wide-Field, Motion-Sensitive Neuron. The Journal of Neuroscience, 19(3), 1122–1141. 10.1523/JNEUROSCI.19-03-01122.1999

Gabbiani, F., Mo, C., & Laurent, G. (2001). Invariance of angular threshold computation in a wide-field looming-sensitive neuron. Journal of Neuroscience, 21(1), 314–329.

Gaten, E., Huston, S. J., Dowse, H. B., & Matheson, T. (2012). Solitary and gregarious locusts differ in circadian rhythmicity of a visual output neuron. Journal of Biological Rhythms, 27(3), 196–205.

Ghalambor, C. K., Angeloni, L. M., & Carroll, S. P. (2010). Behavior as phenotypic plasticity. Evolutionary Behavioral Ecology, 90–107.

Gotham, S., & Song, H. (2013). Non-swarming grasshoppers exhibit density-dependent phenotypic plasticity reminiscent of swarming locusts. Journal of Insect Physiology, 59(11), 1151–1159.

Guo, W., Wu, Z., Yang, L., Cai, Z., Zhao, L., & Zhou, S. (2019). Juvenile hormone–dependent Kazal-type serine protease inhibitor Greglin safeguards insect vitellogenesis and egg production. The FASEB Journal, 33(1), 917–927.

Hawkings, C., & Tamborindeguy, C. (2025). Differential expression of vitellogenin in the brain of Solenopsis invicta workers based on social and nutritional context. Physiological Entomology, 50(1), 64–76. 10.1111/phen.12467

Jang, H., Goodman, D. P., & von Reyn, C. R. (2022). Directional invariance in the Drosophila giant fiber escape circuit. bioRxiv, 2022–07.

Jones, P. W., & Gabbiani, F. (2012). Impact of neural noise on a sensory-motor pathway. J. Neurosci, 32(14), 4923–4934.

Kilpatrick, S. K., Foquet, B., Castellanos, A. A., Gotham, S., Little, D. W., & Song, H. (2019). Revealing hidden density-dependent phenotypic plasticity in sedentary grasshoppers in the genus Schistocerca Stål (Orthoptera: Acrididae: Cyrtacanthacridinae). Journal of Insect Physiology, 118, 103937. 10.1016/j.jinsphys.2019.103937

Kohlmeier, P., Feldmeyer, B., & Foitzik, S. (2018). Vitellogenin-like A–associated shifts in social cue responsiveness regulate behavioral task specialization in an ant. PLOS Biology, 16(6), e2005747. 10.1371/journal.pbio.2005747

Laurent, G., & Davidowitz, H. (1994). Encoding of olfactory information with oscillating neural assemblies. Science, 265(5180), 1872–1875.

Manfredini, F., Brown, M. J. F., & Toth, A. L. (2018). Candidate genes for cooperation and aggression in the social wasp Polistes dominula. Journal of Comparative Physiology A, 204(5), 449–463. 10.1007/s00359-018-1252-6

Marquez-Legorreta, E., Piber, M., & Scott, E. K. (2020). Visual escape in larval zebrafish: Stimuli, circuits, and behavior. In Behavioral and Neural Genetics of Zebrafish (pp. 49–71). Elsevier.

Matheson, T., Rogers, S. M., & Krapp, H. G. (2004). Plasticity in the Visual System Is Correlated With a Change in Lifestyle of Solitarious and Gregarious Locusts. Journal of Neurophysiology, 91(1), 1–12. 10.1152/jn.00795.2003

McMillan, G. A., & Gray, J. R. (2015). Burst firing in a motion-sensitive neural pathway correlates with expansion properties of looming objects that evoke avoidance behaviors. Frontiers in Integrative Neuroscience, 9, 60.

Münch, D., Ihle, K. E., Salmela, H., & Amdam, G. V. (2015). Vitellogenin in the honey bee brain: Atypical localization of a reproductive protein that promotes longevity. Experimental Gerontology, 71, 103–108. 10.1016/j.exger.2015.08.001

Niu, Y., Luo, J., & Zong, C. (2024). Single-cell total-RNA profiling unveils regulatory hubs of transcription factors. Nature Communications, 15(1), 5941. 10.1038/s41467-024-50291-3

O’Shea, M., & Williams, J. L. D. (1974). The anatomy and output connection of a locust visual interneurone; the lobular giant movement detector (LGMD) neurone. Journal of Comparative Physiology, 91(3), 257–266. 10.1007/BF00698057

Ott, S. R., & Rogers, S. M. (2010). Gregarious desert locusts have substantially larger brains with altered proportions compared with the solitarious phase. Proceedings of the Royal Society B: Biological Sciences, 277(1697), 3087–3096. 10.1098/rspb.2010.0694

Panhwar, W. A., & Mustafa, S. B. (2022). Phenotypic plasticity in grasshoppers and locusts: A review. Asian Journal of Science, Engineering and Technology (AJSET), 1(1), 38–51.

Peralta-Rincon, J. R., Escudero, G., & Edelaar, P. (2017). Phenotypic plasticity in color without molt in adult grasshoppers of the genus Sphingonotus (Acrididae: Oedipodinae). Journal of Orthoptera Research, 26, 21–27. 10.3897/jor.26.14550

Peron, S., & Gabbiani, F. (2009). Spike frequency adaptation mediates looming stimulus selectivity in a collision-detecting neuron. Nature Neuroscience, 12(3), 318–326.

Peron, S., Krapp, H. G., & Gabbiani, F. (2007). Influence of electrotonic structure and synaptic mapping on the receptive field properties of a collision-detecting neuron. Journal of Neurophysiology, 97(1), 159–177.

Price, T. D., Qvarnström, A., & Irwin, D. E. (2003). The role of phenotypic plasticity in driving genetic evolution. Proceedings of the Royal Society of London. Series B: Biological Sciences, 270(1523), 1433–1440.

Rind, F. C., & Simmons, P. J. (1999). Seeing what is coming: Building collision-sensitive neurones. Trends in Neurosciences, 22(5), 215–220.

Robinson, M. D., McCarthy, D. J., & Smyth, G. K. (2010). edgeR: a Bioconductor package for differential expression analysis of digital gene expression data. Bioinformatics, 26(1), 139–140.

Rogers, S. M., Harston, G. W. J., Kilburn-Toppin, F., Matheson, T., Burrows, M., Gabbiani, F., & Krapp, H. G. (2010). Spatiotemporal Receptive Field Properties of a Looming-Sensitive Neuron in Solitarious and Gregarious Phases of the Desert Locust. Journal of Neurophysiology, 103(2), 779–792. 10.1152/jn.00855.2009

Rogers, S. M., Krapp, H. G., Burrows, M., & Matheson, T. (2007). Compensatory Plasticity at an Identified Synapse Tunes a Visuomotor Pathway. The Journal of Neuroscience, 27(17), 4621–4633. 10.1523/JNEUROSCI.4615-06.2007

Rogers, S. M., Riley, J., Brighton, C., Sutton, G. P., Cullen, D. A., & Burrows, M. (2016). Increased muscular volume and cuticular specialisations enhance jump velocity in solitarious compared with gregarious desert locusts, Schistocerca gregaria. Journal of Experimental Biology, 219(5), 635–648. 10.1242/jeb.134445

Schlotterer, G. (1977). Response of the locust descending movement detector neuron to rapidly approaching and withdrawing visual stimuli. Canadian Journal of Zoology, 55(8), 1372–1376.

Simões, P. M. V., Ott, S. R., & Niven, J. E. (2016). Environmental Adaptation, Phenotypic Plasticity, and Associative Learning in Insects: The Desert Locust as a Case Study. Integrative and Comparative Biology, 56(5), 914–924. 10.1093/icb/icw100

Smith, T., Heger, A., & Sudbery, I. (2017). UMI-tools: Modeling sequencing errors in Unique Molecular Identifiers to improve quantification accuracy. Genome Research, 27(3), 491–499.

Song, H., Foquet, B., Mariño-Pérez, R., & Woller, D. A. (2017). Phylogeny of locusts and grasshoppers reveals complex evolution of density-dependent phenotypic plasticity. Scientific Reports, 7(1), 6606. 10.1038/s41598-017-07105-y

Sun, H., & Frost, B. J. (1998). Computation of different optical variables of looming objects in pigeon nucleus rotundus neurons. Nature Neuroscience, 1(4), 296–303. 10.1038/1110

Sword, G. A. (2002). A role for phenotypic plasticity in the evolution of aposematism. Proceedings of the Royal Society of London. Series B: Biological Sciences, 269(1501), 1639–1644.

Sword, G. A. (2003). To be or not to be a locust? A comparative analysis of behavioral phase change in nymphs of Schistocerca americana and S. gregaria. Journal of Insect Physiology, 49(7), 709–717.

Temizer, I., Donovan, J. C., Baier, H., & Semmelhack, J. L. (2015). A visual pathway for looming-evoked escape in larval zebrafish. Current Biology, 25(14), 1823–1834.

Thompson, T. (1999). Genotype–environment interaction and the ontogeny of diet-induced phenotypic plasticity in size and shape of Melanoplus femurrubrum (Orthoptera: Acrididae). Journal of Evolutionary Biology, 12(1), 38–48.

Uvarov, B. (1966). British Grasshoppers and Crickets. Nature, 210(5031), 63–64.

Von Reyn, C., & Card, G. (2012). The role of the giant fibers in visually evoked escape behavior. Front. Behav. Neurosci. Conference Abstract: Tenth International Congress of Neuroethology.

Whitman, D. W., & Agrawal, A. A. (2009). What is phenotypic plasticity and why is it important. Phenotypic Plasticity of Insects: Mechanisms and Consequences, 1–63.

Xiao, Q., Li, D.-P., & Wang, S.-R. (2006). Looming-sensitive responses and receptive field organization of telencephalic neurons in the pigeon. Brain Research Bulletin, 68(5), 322–328.

Zhu, Y., & Gabbiani, F. (2016). Fine and distributed subcellular retinotopy of excitatory inputs to the dendritic tree of a collision-detecting neuron. Journal of Neurophysiology, 115(6), 3101–3112.

Zhu, Y., & Gabbiani, F. (2018). Combined Two-Photon Calcium Imaging and Single-Ommatidium Visual Stimulation to Study Fine-Scale Retinotopy in Insects. In R. V. Sillitoe (Ed.), Extracellular Recording Approaches (Vol. 134, pp. 185–206). Springer New York. 10.1007/978-1-4939-7549-5_10

